# Chromatin states contribute to coordinated allelic transcriptional bursting to drive iPSC reprogramming

**DOI:** 10.1101/2023.07.13.548864

**Authors:** A Parichitran, HC Naik, AJ Naskar, LS Bammidi, S Gayen

## Abstract

Molecular mechanisms behind the reprogramming of somatic cells to induced pluripotent stem cells (iPSC) remain poorly understood. While dynamic changes in gene expression are considered to drive reprogramming, the contribution of individual alleles of genes to reprogramming remains unexplored. It is thought that two alleles of a gene can transcribe independently or coordinatedly, which in turn can lead to temporal expression heterogeneity with potentially distinct impacts on cell fate. Here, we profiled genome-wide transcriptional burst kinetics with an allelic resolution during the reprogramming of mouse embryonic fibroblast (MEF) to iPSC. We show that many genes involved in iPSC reprogramming pathways exhibit bursty expression and contribute to dynamic autosomal random monoallelic expression (aRME). Moreover, we find that the degree of coordination of allelic bursting differs among genes and changes dynamically during iPSC reprogramming. Importantly, we show that alleles of many reprogramming-related genes burst in a highly coordinated fashion. ATAC-seq analysis revealed that coordination of allelic bursting is linked to allelic chromatin accessibility. Consistently, we show that highly coordinated genes are enriched with chromatin accessibility regulators such as H3K36me3, H3K27ac, histone variant H3.3 and BRD4. Collectively, our study demonstrates that chromatin states contribute to coordinated allelic bursting to fine-tune the expression of genes involved in iPSC reprogramming and provides insights into the implications of allelic bursting coordination in cell fate specification.

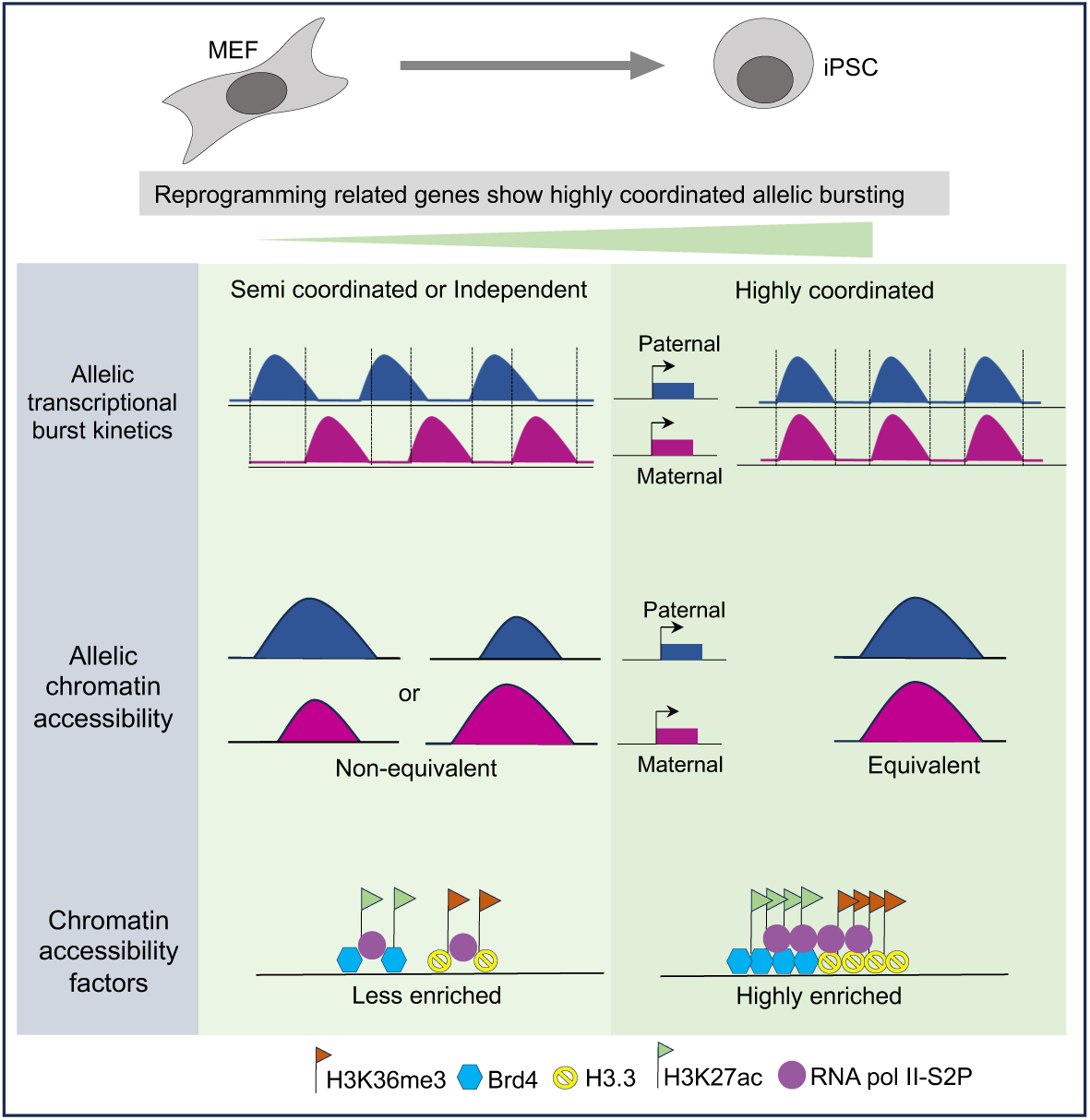

## Introduction

Reprogramming of somatic cells to induced pluripotent stem cells (iPSC) serves as an excellent model system for studying the mechanisms of cell fate specification and gene regulation (Bauer et al., 2021; Cheloufi et al., 2015; Generoso et al., 2023; Naik et al., 2023; Onder et al., 2012; Polo et al., 2012; Takahashi and Yamanaka, 2006, 2016). However, mechanisms of iPSC reprogramming remain poorly understood. Precise expression of genes involved in iPSC reprogramming is crucial for reprogramming. To date, the gene expression dynamics during iPSC reprogramming have been studied at the gene level but not at the allelic level. In eukaryotic cells, transcription happens in a sporadic manner through random transcriptional bursting separated by periods of silent state, which contributes to the gene expression heterogeneity among the identical cells (Larsson et al., 2019; Little et al., 2013; Naik et al., 2021; Padovan-Merhar et al., 2015; Raj and van Oudenaarden, 2008; Raj et al., 2006; RV et al., 2021). On the other hand, the regulation of transcriptional bursting can be shared or autonomous between alleles (Finn et al., 2019; Naik et al., 2021; Onuchic et al., 2018; RV et al., 2021). Indeed, recent studies by us and others demonstrated that often the kinetics of transcriptional bursting of the two alleles of a gene differs and which in turn gives rise to dynamic random monoallelic expression (dynamic RME) (Borel et al., 2015; Gendrel et al., 2016; Gregg, 2017; Naik et al., 2021; Reinius and Sandberg, 2015; Reinius et al., 2016; RV et al., 2021). It is believed that dynamic RME can induce temporal variations of gene expression among cells and thereby may contribute to cell fate specification. Therefore, it is important to profile the transcriptional kinetics of genes at the allelic level to understand how gene expression is fine-tuned for precise cell fate specification. However, allelic transcriptional kinetics during iPSC reprogramming remain unexplored. To address this, we have profiled genome-wide transcriptional burst kinetics at allelic level across different stages of reprogramming of mouse embryonic fibroblast (MEF) to iPSC using allele-specific single-cell RNA-Sequencing (scRNA-seq) analysis. We find that the degree of coordination of allelic bursting differs among genes and changes dynamically during iPSC reprogramming. Importantly, we find that many genes involved in reprogramming pathways have a high degree of allelic coordination. On the other hand, the factors involved in coordinating allelic bursting are not known. Transcription factors and enhancer functions are thought to modulate transcriptional burst kinetics (Larsson et al., 2019). Emerging trends suggest that chromatin states are linked to burst kinetics (Chen et al., 2019; Fraser et al., 2021; Nicolas et al., 2018). Therefore, we have explored how chromatin states contribute to the coordinated allelic transcriptional bursting during the iPSC reprogramming.

## Result

### Prevalent bursty expression contributes to dynamic aRME during iPSC reprogramming

Transcription of many genes occurs in a stochastic manner, where genes undergo sporadic bursting to produce RNA. The kinetics of transcriptional bursting is deduced through the well-known “two-state model” of transcription. According to the “two-state model,” the promoter of a gene switches stochastically from an inactive/OFF state to an active/ON state and burst kinetics is determined through two parameters: burst frequency and burst size (Fig 1A) (Chubb et al., 2006; Larson, 2011; Raj and van Oudenaarden, 2008). Burst frequency is defined by the number of bursts per unit time and burst size explains the mean amount of mRNA molecules when the gene is in active state (Fig 1A). To date, burst kinetics have been studied mainly at the gene level, not the allelic level. However, profiling burst kinetics at allelic resolution is important as the kinetics of bursting between alleles often differ (Naik et al., 2021). Here, we have delineated the kinetics of allelic bursting genome-wide across different stages of reprogramming of MEF to iPSC (Fig 1B). To profile, genome-wide allelic transcriptional burst kinetics, we performed SCALE (Single-Cell ALlelic Expression) analysis using scRNA-seq datasets (Fig 1B). SCALE relies on Empirical Bayes Framework, which first classifies the genes into: monoallelic, biallelic and silent based on the allele-specific read counts and the biallelic genes are further categorized into: biallelic bursty and biallelic non-bursty (Fig 1C) (Jiang et al., 2017). The MEF cells used in the experiment were derived from a cross of two divergent mouse strains *M. Musculus* (129S1) and *M. Castaneous* (CAST) and thereby enabling us to perform allele-specific analysis based on strain-specific SNPs (Fig 1B) (Talon et al., 2021). We excluded low-expressed genes from our study to avoid allelic dropout-related technical noise, which could lead to inaccurate estimation of allelic expression (Kim et al., 2015; Santoni et al., 2017; Zhao et al., 2017). Through SCALE analysis, we found that most of the biallelic genes (∼90%) have bursty expression across all stages of reprogramming (Fig 1D). Next, we compared the bursty/non-bursty pattern of gene expression across different reprogramming stages and found that most genes (n=777) maintained bursty expression throughout the reprogramming (Fig 1E). Interestingly, these genes were enriched toward key reprogramming-related processes, including blastocyst growth (Fig 1F). Moreover, genes (n=105) that remained bursty on all days except iPSCs were also enriched in many reprogramming-related biological processes, including chordate embryonic development (Fig 1F). These results indicated that many genes involved in iPSC reprogramming pathways exhibit bursty expression. However, a few genes (n=22) that maintained bursty expression in all day points except MEF did not show enrichment towards iPSC reprogramming pathways (Fig S1A). Genes (n=17), which remained non-bursty throughout the reprogramming, were not enriched to reprogramming pathways (Fig S1A). On the other hand, we have previously shown that bursty expression results in dynamic RME, which creates gene expression heterogeneity among cells (Naik et al. 2021). Therefore, we explored the landscape of allelic expression throughout the different stages of iPSC reprogramming by allele-specific scRNA-seq analysis. We categorized a gene as monoallelic within a cell if at least 95% of the expression originated from one allele. We classified the monoallelic expression into three categories: nonrandom monoallelic, random monoallelic with one allele and random monoallelic with either allele. Our analysis revealed that ∼80-90% of genes have dynamic RME across different stages of reprogramming (Fig 1G). Analysis of allelic expression of X-linked genes in MEF showed monoallelic expression from the CAST allele as 129S1-X is inactivated in these cells, thereby validating our allele-specific analysis pipeline (Fig S1B). X-inactivation is a process through which female mammals compensate the dosage of X-linked gene expression between sexes (Gayen et al., 2015, 2016; Kaur et al., 2019; Saiba et al., 2018; Samanta et al., 2022; Sarkar et al., 2015). Altogether, our analysis suggested widespread bursty expression of genes during iPSC reprogramming, resulting in dynamic aRME.

**Figure 1:**
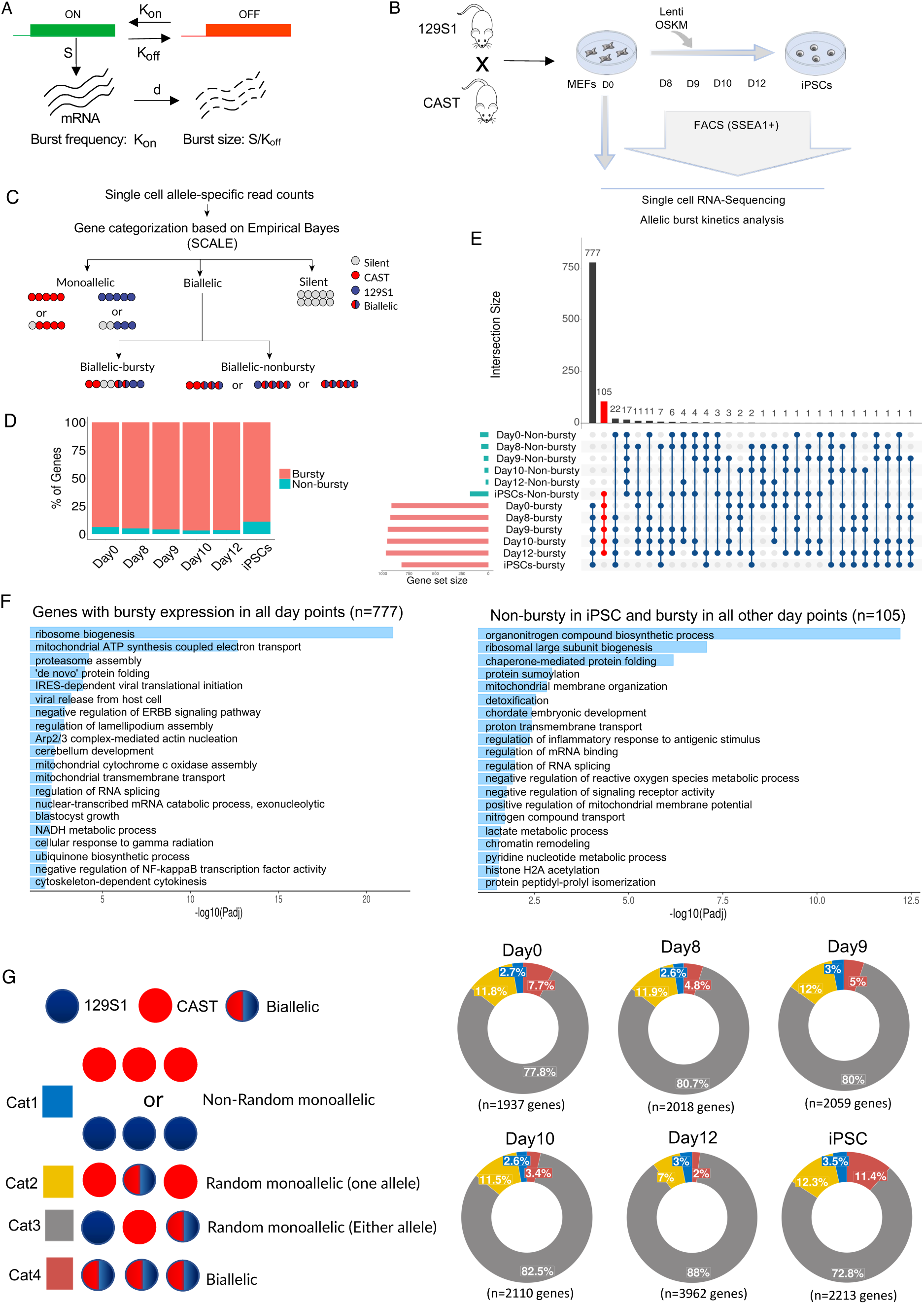
Genome-wide profiling of bursty expression and dynamic aRME in different stages of MEF to iPSC reprogramming. **(A)** Diagrammatical representation of the “two-state model” of transcriptional bursting. K_on_ is the rate of transcriptional activation of gene; Koff is the rate at which a gene becomes transcriptionally inactive; S is the rate of transcription of a gene in active state; d is the rate of mRNA decay; burst kinetics is represented through K_on_ (burst frequency) and S/K_off_ (burst size). (B) Graphical representation of OSKM (Oct-3/4, Sox2, Klf4 and c-Myc) mediated reprogramming of hybrid MEF cells (129S1 X CAST) and collection of cells of intermediate stages day 8, day 9, day 10, day 12 and iPSCs. (C) Outline of categorization of bursty and non-bursty gene through SCALE analysis. (D) Quantification of the extent of bursty genes across all day points (MEF, day 8, day 9, day 10, day 12, iPSC) of reprogramming. (E) Cross comparison plot of bursty and non-bursty genes across the six stages of reprogramming. (F) Gene ontology (GO) enrichment analysis of 777 genes that remain bursty across all day points and 105 genes that are non-bursty in iPSCs but bursty in other day points. (G) Plots representing the percent of genes with different allelic expression categories throughout different stages of reprogramming; Cat 1: non-random monoallelic, Cat 2: random monoallelic with one allele, Cat 3: random monoallelic with either allele, Cat 4: biallelic.

### Alleles of genes exhibit similar burst kinetics but have different coordination during iPSC reprogramming

Next, we investigated if the two alleles of a gene have similar burst kinetics or not to better understand the bursty gene expression during iPSC reprogramming. To explore this, we profiled burst frequency and burst size at the allelic level for the biallelic bursty genes. We observed a high degree of correlation of both burst frequency (r= 0.64-0.778) and burst size (r= 0.731-0.772) between two alleles across all day points (Figure 2A and 2B). Very few genes exhibited significant burst frequency and size differences between the two alleles, as marked by the red triangles (Figure 2A and 2B). Taken together, our results suggested that alleles of most of the genes have similar burst kinetics. Next, we explored the degree of coordination of bursting between two alleles by plotting the percent of cells expressing neither allele (p_0_) vs. the percent of cells expressing both alleles (p_2_) (Figure 2C). We broadly categorized the degree of coordination of two alleles’ bursting into three categories: (1) Highly coordinated genes between the two dotted blue diagonal lines (gray asterisks). (2) Independent genes that lie between the uppermost and lowermost red dotted curve lines (rosewood triangles) and (3) Semi-coordinated genes that lie between the uppermost red curve line and lower blue dotted diagonal line (Persian blue dots) (Figure 2C). We observed that the majority of genes exhibited semi-coordinated allelic bursting at all-day points. However, many genes also showed highly coordinated allelic bursting across the different stages of reprogramming (Figure 2C).

**Figure 2:**
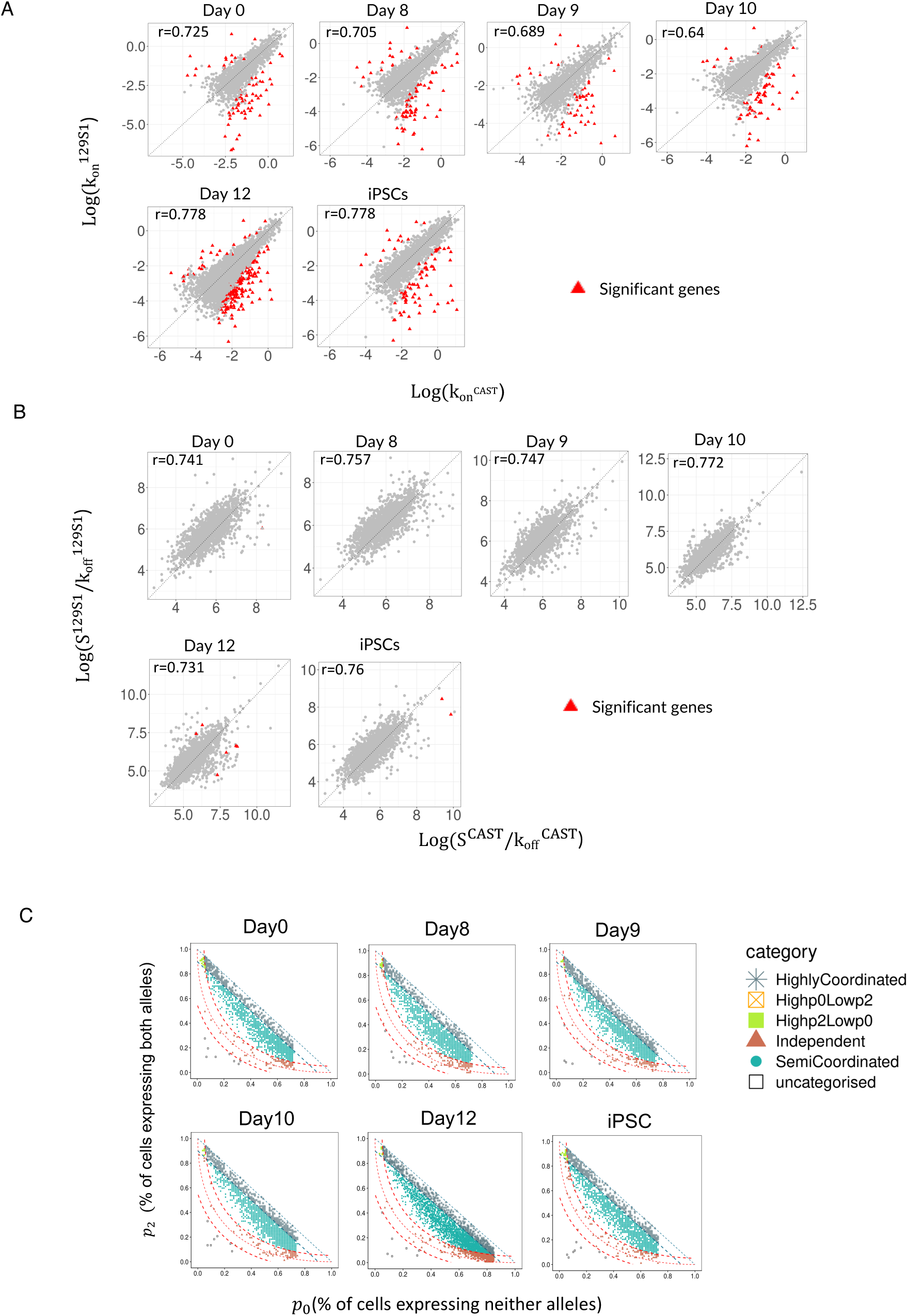
Profiling of allelic burst kinetics and coordination. Plots representing correlation between (A) allelic burst frequency across different stages of reprogramming day 0 MEF (r = 0.725), day 8 (r = 0.705), day 9 (r = 0.689), day 10 (r =0.64), day 12 (r = 0.778) and iPSCs (r = 0.778) and (B) allelic burst size in day 0 MEF (r= 0.741), day 8 (r= 0.757), day 9 (r= 0.747), day 10 (r=0.772), day 12 (r= 0.731) and iPSCs (r= 0.76). Genes that exhibit significant difference in burst frequency and size between two alleles have been marked by red triangles. (C) Smooth scatterplots representing bursting coordination between two alleles of genes for day 0 MEF, day 8, day 9, day 10, day 12 and iPSCs. Percent of cells expressing neither allele (p0) is plotted with percent of cells expressing both alleles (p2), blue diagonal line represents perfect coordination (p0+p2=1), while red curve signified independent bursting with shared kinetics. Different categories of genes based on allelic bursting coordination: low p0 high p2 (green filled squares), high p0 and low p2 (marked by orange unfilled squares), perfectly coordinated (p0+p2>0.90 marked by gray asterisk between blue dotted diagonal lines), independent genes marked by rosewood triangles (between upper and lower red curved lines, with threshold of +0.05 signified by upper red curve and -0.05, signified by lower red curve) and semi-coordinated genes marked with persian blue dots.

### Genes involved in iPSC reprogramming undergo highly coordinated allelic bursting

Next, we performed a gene ontology (GO) enrichment analysis of highly coordinated genes across all stages of reprogramming (Figure 3). Interestingly, we found that highly coordinated genes in iPSCs are enriched in processes linked to iPSCs like stem cell population maintenance, in-utero embryo development, endoderm formation etc. (Figure 3A). Furthermore, highly coordinated genes in iPSCs enriched in cellular respiration, cristae formation and glutamine metabolism, which are relevant to metabolic remodeling in iPSCs (Figure 3A) (Mathieu and Ruohola-Baker, 2017; Teslaa and Teitell, 2015; Tohyama et al., 2017). Importantly, genes involved in ribosome biogenesis, crucial for stem cell maintenance, were highly enriched into the highly coordinated gene cohort in iPSC (Figure 3A) (Gabut et al., 2020). On the contrary, highly coordinated genes in Day0-MEF cells were not enriched in such stem cell-related pathways. However, across the different intermediate stages of reprogramming, highly coordinated genes were enriched towards many reprogramming-related biological processes such as ribosome biogenesis, protein folding, stem cell differentiation, NFkB signaling etc. (Figure 3B) (Gabut et al., 2020; Kaltschmidt et al., 2021; Yan et al., 2020). Next, we performed a cross-comparison of the three allelic coordination categories across all-day points and found that while many genes maintained a similar degree of allelic coordination across all-day points, some genes did not (Figure 3C). Strikingly, we found that the genes (n=59) that become highly coordinated in iPSCs enriched towards ribosome biogenesis, aerobic respiration, cristae formation, glycine metabolism, mitochondrial respiratory chain assembly and endoderm formation all of which are highly critical and directly relevant for iPSC reprogramming (Figure 3D) (Kang et al., 2019; Teslaa and Teitell, 2015; Xu et al., 2013). Moreover, genes that remain highly coordinated (n=38) on all days showed enrichment towards some iPSC reprogramming linked functions like protein folding, protein stability, NFkB pathway, mitochondrial electron transport etc. (Figure 3D). Altogether, our analysis revealed that the two alleles of many genes involved in iPSC reprogramming have a high degree of transcriptional bursting coordination. In parallel, we also observed that genes that remained semi-coordinated (n=318) on all days or converted from highly coordinated on day 0 to semi-coordinated on other days (n=40) enriched towards some iPSC reprogramming linked functions like proteasome assembly, negative regulation of NFkB pathway, oxidative stress response etc. (Figure S2) (Hawkins et al., 2016; Mathieu and Ruohola-Baker, 2017; Schröter and Adjaye, 2014; Szutorisz et al.).

**Figure 3:**
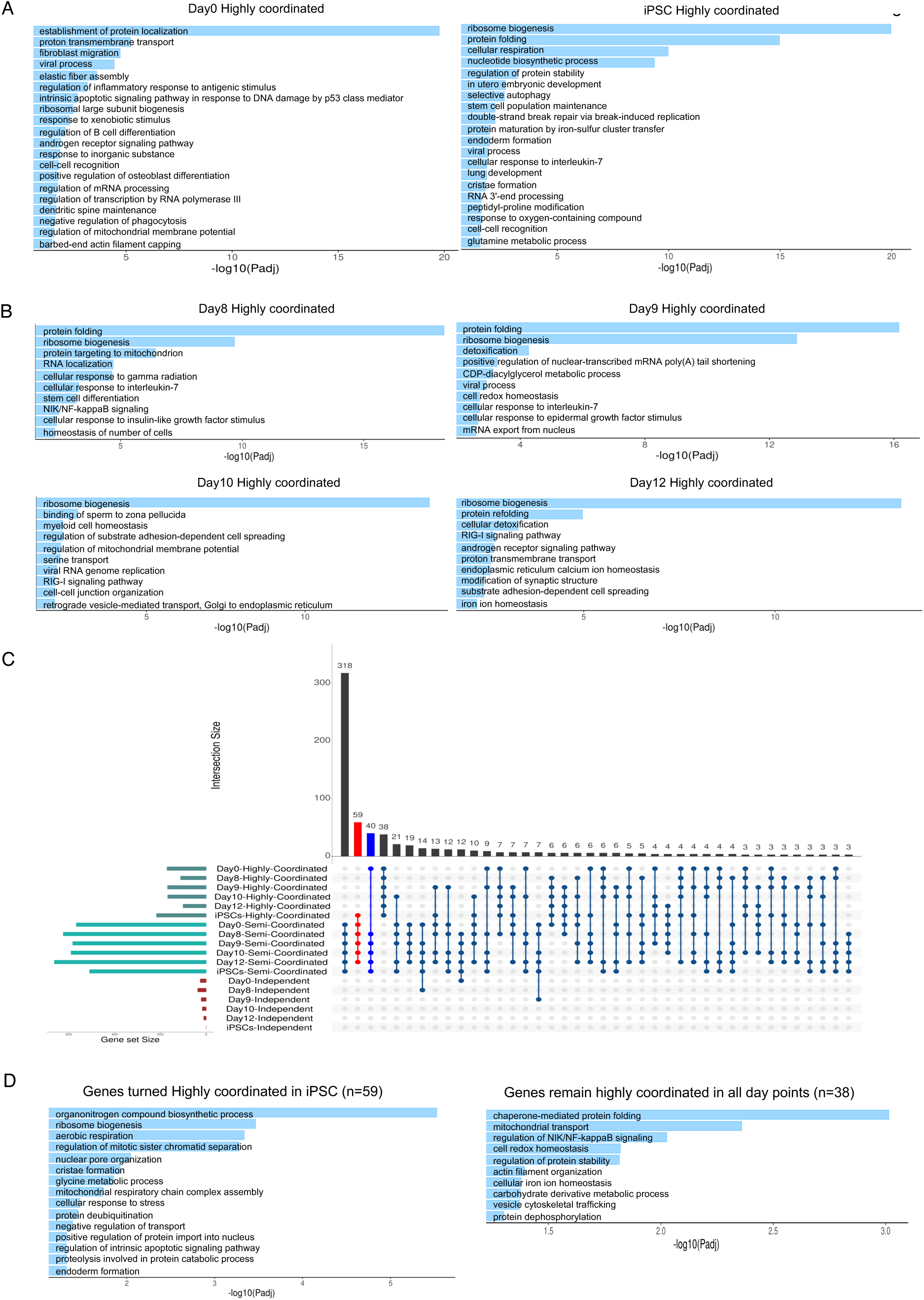
Highly coordinated genes are enriched to iPSC reprogramming related processes. (A) Gene ontology (GO) enrichment analysis of highly coordinated genes in day 0 MEF and iPSCs. (B) Gene ontology (GO) enrichment analysis of highly coordinated genes in intermediate stages of reprogramming: day 8, day 9, day 10 and day 12 cells. (C) Cross comparison plot of highly coordinated, semi-coordinated and independent genes across day 0 MEF, day 8, day 10, day 12 and iPSCs. (D) Gene ontology (GO) enrichment analysis of genes (n=59) that become highly coordinated in iPSCs (left) and genes (n=38) that maintained highly coordinated allelic bursting through all day points.

### Coordinated allelic bursting is linked to chromatin accessibility

To understand the mechanisms of allelic bursting coordination, we asked whether the degree of coordination of allelic bursting is linked to the chromatin states. To address this, we profiled genome-wide allelic chromatin accessibility across different stages of MEF to iPSC reprogramming through allele-specific analysis of available ATAC-sequencing (ATAC-seq) datasets (Talon et al., 2021). The same hybrid MEFs (129S1 X CAST), as described for scRNA-seq, were used for this experiment, allowing us to profile chromatin accessibility at the allelic level. We analyzed ATAC-seq in MEFs (day 0) and across reprogramming stages (SSEA1+ reprogramming intermediates at days 8, 9, 10, 12 and iPSCs), like the burst kinetics analysis (Fig 4A). We first validated our allele-specific ATAC-seq analysis pipeline by quantifying the difference in the enrichment of ATAC-seq reads between active-X (129S1) vs. inactive-X (CAST allele) (Fig S3). In consistence with previous reports, in MEFs and early reprogramming intermediates, active-X (CAST) showed strong enrichment of ATAC-seq reads, whereas the inactive-X (129S1) showed almost no enrichment and upon reactivation of the inactive-X towards the attainment of iPSCs, there was a gain of chromatin accessibility (Fig S3). Taken together, enrichment analysis of ATAC-seq reads of X-linked genes validated the accuracy of our method. Next, we compared the enrichment of ATAC-seq reads between two alleles of different categories of genes (highly coordinated, semi-coordinated and independent) across 3 kb upstream and 3 kb downstream of TSS during reprogramming (Fig 4A). Interestingly, our analysis revealed that the two alleles of highly coordinated genes have very similar enrichment in most day points. Whereas enrichment of ATAC-seq reads of allele of semi-coordinated/independent genes differed in most cases (Fig 4A). As expected, allelic enrichment of ATAC-seq reads considering all autosomal genes was quite similar (Fig 4A). Together, our analysis suggested a positive correlation between the coordination of allelic bursting and the similarity of allelic chromatin accessibility. Furthermore, we find a similar trend for genes (n=59), which switches from a semi-coordinated state to highly coordinated upon attainment of iPSC (Fig 4B). On the other hand, genes that maintained highly coordinated or semi-coordinated bursting throughout reprogramming and switched from highly coordinated in MEF to semi-coordinated on other days showed almost similar allelic chromatin accessibility (Data not shown).

**Figure 4:**
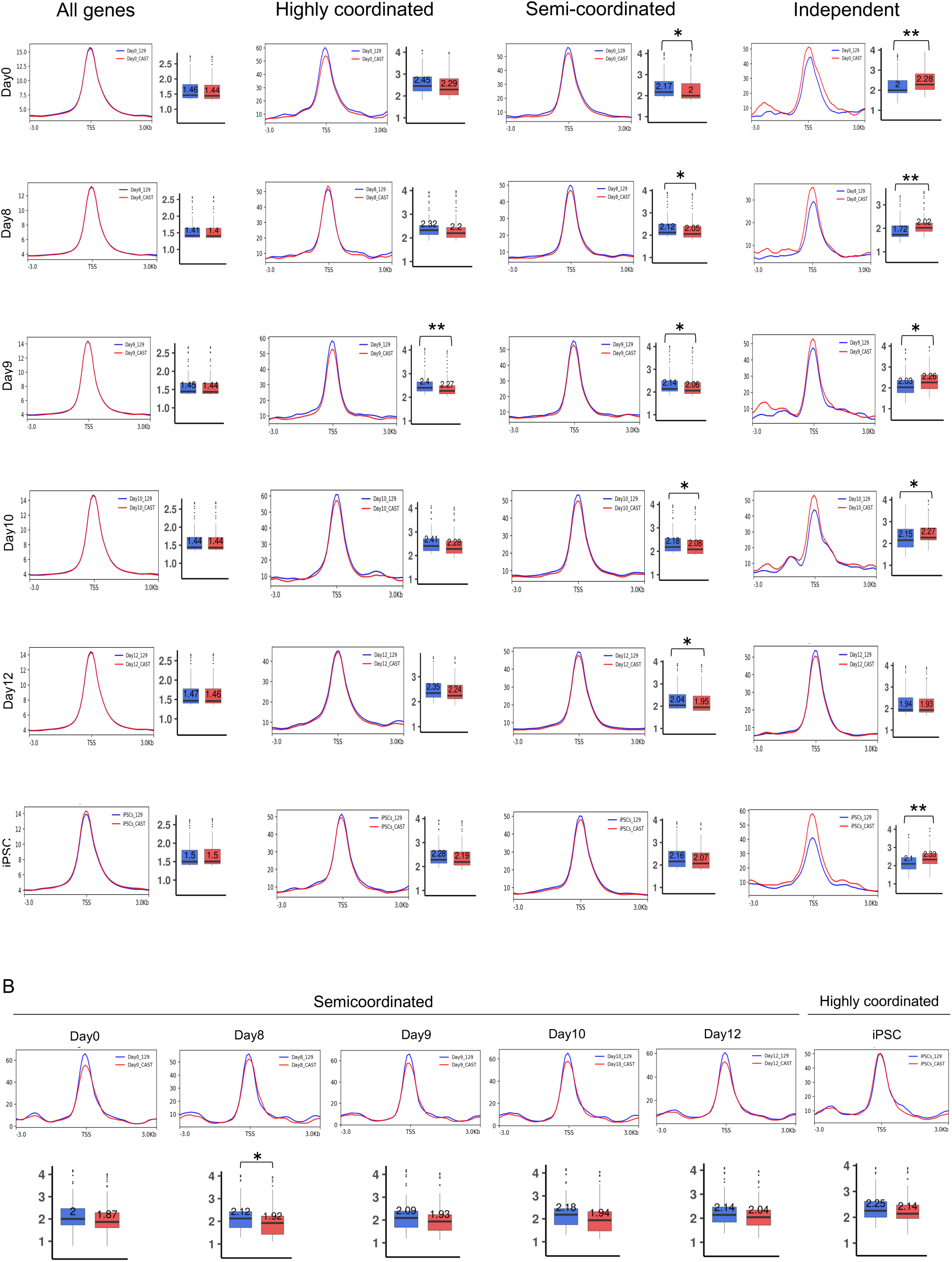
Comparison of allelic chromatin accessibility of highly coordinated, semi-coordinated and independent genes. (A) Quantitative analysis of allelic accessibility enrichment across 3 kb upstream and 3 kb downstream of transcription start-site (TSS) of all autosomal genes, highly coordinated, semi-coordinated and independent genes throughout different stages of reprogramming: reprogramming: day 0, day 8, day 10, day 12 and iPSCs. In the boxplots, the line inside each of the boxes denote median value and edges of each box represent 25% and 75% of dataset respectively (Mann-Whitney U Test: p *value < 0.01*; ** and p *value < 0.05*; *) (B) Plots representing allelic accessibility enrichment for genes that converted from semi-coordinated in all days to highly coordinated in iPSCs (n=59). In the boxplots, the line inside each of the boxes denote median value and edges of each box represent 25% and 75% of dataset respectively (Mann-Whitney U Test: p *value < 0.01*; ** and p *value < 0.05*; *)

### Chromatin accessibility factors contribute to allelic bursting coordination

Next, we investigated if the degree of allelic coordination is dependent on chromatin accessibility factors enrichment. To explore this, we profiled the enrichment of different chromatin accessibility factors in the TSS and gene body of highly coordinated, semi-coordinated and independent genes in MEF and iPSCs. Interestingly, we found that H3K36me3, H3K27ac, H3.3 and RNA_PolII-S2P are highly enriched on the TSS/gene body of highly coordinated genes compared to the semi-coordinated / independent genes in both MEF and iPSC (Fig 5A). Moreover, in iPSC, we found that highly coordinated genes are enriched with chromatin remodeler BRD4 (Fig 5A). H3K79me2, H3K9ac, and RNA PolII-S5P were highly enriched to the highly coordinated genes in MEF cells (Fig 5A). Next, we tested if switching of coordination pattern of genes in MEF to iPSC is associated with changes in the pattern of chromatin accessibility factors enrichment. We found that highly coordinated genes in MEF having higher enrichment of H3K36me3, H3K27ac, H3.3 and RNA_PolII-S2P compared to the semicordited genes flips their pattern of enrichment in iPSC upon conversion of their coordination pattern (Fig 5B). Whereas genes that do not switch their coordination pattern in MEF to iPSC maintain the enrichment pattern of these marks (Fig 5B). Altogether, our analysis suggested that higher enrichment of these chromatin accessibility factors ensures highly coordinated allelic bursting. However, other chromatin modifications, such as H3K9/14ac, H3K4me1, H3K4me2, H3K4me3, H3K9me3 and H3K27me3, did not show such differences in enrichment between different categories of genes (Fig S4).

**Figure 5:**
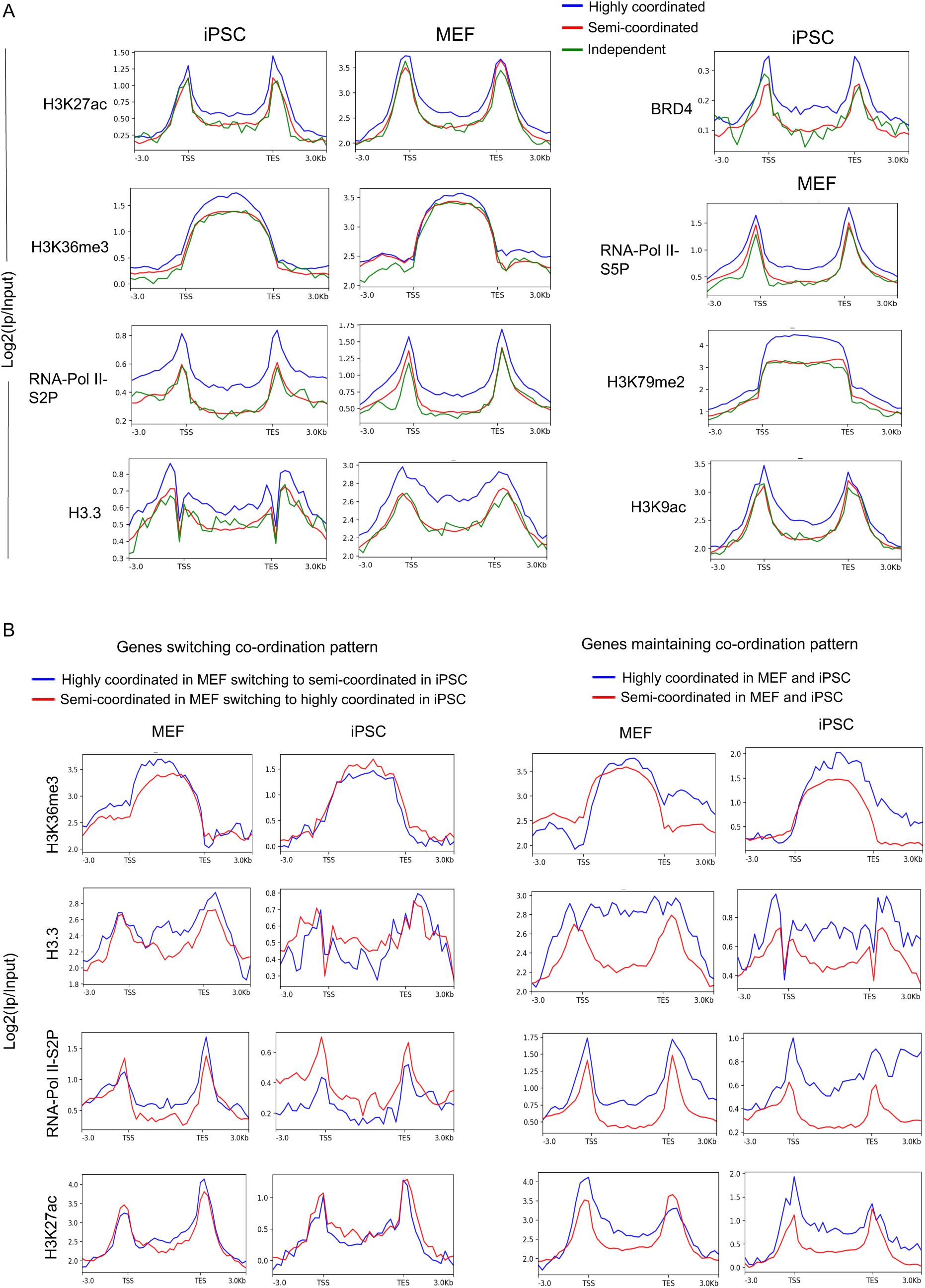
Correlation between occupancy of chromatin marks and degree of allelic bursting coordination: (A) Comparison of enrichment of different chromatin accessibility related factors (H3K27ac, H3K36me3, H3K9ac, H3K79me2, H3.3, BRD4, RNA Pol-II-S2P and RNA Pol-II-S5P) in the gene body and TSS of highly coordinated, semi-coordinated and independent genes in MEF and iPSC. (B) Plots representing quantitative enrichment of different chromatin accessibility related factors (H3K27ac, H3.3, H3K36me3 and RNA Pol-II-S2P) of the gene body and TSS of genes switching or maintaining coordination pattern between MEF and iPSCs.

## Discussion

Since the discovery of reprogramming of somatic cells to pluripotent state in 2006, the underlying precise mechanistic aspect of reprogramming remains unknown (Takahashi and Yamanaka, 2006). To fill this gap, it is imperative to understand the transcriptional regulation of reprogramming related genes quantitatively. Emerging trends suggest that transcriptional regulation of two alleles of a gene is not always shared and can be independent and which in turn can lead to temporal expression heterogeneity (Finn et al., 2019; Naik et al., 2021). Therefore, it is important to explore the allelic contribution of genes and their cooperativity to understand how cells fine-tune the optimal expression to bring developmental precision. To understand this aspect quantitatively, we have profiled genome-wide transcriptional burst kinetics at the allelic level and their relevance to cell state transition during iPSC reprogramming. We found that most of the autosomal genes exhibit bursty expression and have dynamic aRME (∼73-88%) across different stages of iPSC reprogramming (Fig 1), which is consistent with our previous reports in pre-gastrulation embryos (Naik et al., 2021). Importantly, we found that many genes involved in iPSC reprogramming pathways exhibit bursty expression (Fig 1). Interestingly, we found that burst frequency and burst size are highly similar between two alleles for most of the genes across reprogramming (Fig 2A and 2B). However, in terms of cooperativity of allelic bursting, we found different patterns of allelic bursting: while most of the genes exhibited semi-coordinated allelic bursting, many genes showed highly coordinated allelic bursting (Fig 2C). On the other hand, few genes showed independent nature of allelic bursting.

Next, we found that the degree of coordination of allelic transcriptional bursting is relevant to reprogramming pathways. We show that allelic bursting of many genes crucial to iPSC reprogramming occurs in a highly coordinated fashion (Fig 3). We found that genes related to translation, protein stability, protein folding and RNA processing undergo highly coordinated allelic bursting in iPSC. Specially, ribosome biogenesis-related genes (e.g., *rpl7, nop58*, *nmd3, nifk* etc.) becomes highly coordinated upon initiation of reprogramming (d8 onwards) and remained highly coordinated through intermediate stages and iPSC (Fig 3). Indeed, reprogramming of ribosome biogenesis is crucial for iPSC reprogramming and stem cell maintenance to enhance translational efficiency (Gabut et al., 2020; Hu, 2020). Moreover, pluripotent embryonic stem cells bear a high density of inactive ribosomes to facilitate increased translation efficiency during their subsequent differentiation into different lineages (Gabut et al., 2020; Novak et al., 2012; Sampath et al., 2008; You et al., 2015). Additionally, important genes like *nanog*, *dppa2, med28 etc.,* which play important roles in iPSC reprogramming, showed highly coordinated allelic expression (Li et al., 2015). Importantly, many genes related to embryonic development and stem cell maintenance showed highly coordinated allelic bursting (Fig 3). Surprisingly, we observed that genes associated with the mitochondrial cristae formation and mitochondrial respiratory chain assembly (ndufa10, coa3 and samm50) exhibited highly coordinated allelic bursting in iPSC (Fig 3). It is known that iPSC reprogramming is associated with the metabolic shift from oxidative phosphorylation (OXPHOS) to glycolysis, which is accompanied by the formation of immature spherical mitochondria with less dense cristae in iPSCs in comparison to the highly elongated mitochondria with dense cristae network in MEF (Seo et al., 2018; Teslaa and Teitell, 2015; Xu et al., 2013). Although reprogramming to iPSC is associated with the shift from OXPHOS to glycolysis, iPSC remains dependent on mitochondrial metabolism for intermediate metabolites, which plays important role in epigenomic regulation to drive iPSC reprogramming or stem cell pluripotency (Carey et al., 2014; Fang et al., 2019; Guitart et al., 2017; Moussaieff et al., 2015; Todd et al., 2010; Zhang et al., 2016, 2018; Zhou et al., 2016). Therefore, it may be possible that highly coordinated bursting of these mitochondria-related genes is crucial for iPSC reprogramming. Notably, genes involved in nuclear pore formation shifted to highly coordinated state in iPSC. It has been demonstrated that the nuclear pore complex plays an important role in modulating pluripotency and reprogramming (Hansson et al., 2012; Yang et al., 2014). On the other hand, in a recent study, we demonstrated that developmental genes related to gastrulation undergo highly coordinated allelic bursting in pre-gastrulation mouse embryos (Naik et al., 2021). Taken together, we conclude that many genes crucial to reprogramming and development burst in a highly coordinated fashion. Possibly, the coordination of transcriptional bursting between the two alleles fine-tune gene expression dosage to drive precise development or reprogramming.

Next, we show that chromatin states contribute to the coordination between allelic transcriptional bursting. We demonstrate that the coordination of allelic bursting is linked to the open chromatin. We find that with the increased degree of allelic coordination of bursting, coordination of allelic chromatin accessibility also increases (Fig 4). Previous studies have shown that random chromatin accessibility contributes to transcriptional bursting by providing intermittent accessibility to the transcription factors (Brown et al., 2013; Chen et al., 2019; Fraser et al., 2021; Nicolas et al., 2018). Moreover, it is believed that open chromatin states modulate transcriptional burst size and burst frequency (Bartman et al., 2016; Elise Bullock et al., 2022; Fraser et al., 2021; Nicolas et al., 2018). Our analysis extends the support on the role of chromatin accessibility on transcriptional bursting and, most importantly, sheds light on how allelic open chromatin dynamics is linked to allelic transcriptional burst kinetics. In the future, more extensive studies would help to gain deeper insights into the role of chromatin accessibility in mediating allelic bursting coordination.

On the other hand, we find that highly coordinated genes are enriched with important chromatin accessibility factors: H3K36me3, H3K27ac, BRD4 and histone variant H3.3 (Fig 5A). In fact, H3K36me3, H3K27ac, along with BRD4, have been reported to play a role in orchestrating transcription and burst frequency (Abe et al., 2022; Altendorfer et al., 2022; Nicolas et al., 2018; Ochiai et al., 2020; Pal et al., 2023; Sundarraj, 2021). Interestingly, we report the implications of H3.3 in allelic transcriptional bursting for the first time. Moreover, we found that H3K79me2, H3K9ac, RNA PolII-S2P and RNA PolII-S5P were also highly enriched in highly coordinated genes. Taken together, we propose that these chromatin-related factors play a crucial role in mediating the coordination of allelic bursting. In the future, analysis of allele-specific enrichment of these marks or factors in highly coordinated, semi-coordinated and independent genes would provide better insight into the plausible regulatory link among allele-specific enrichment of active chromatin marks, burst frequency and degree of coordination of bursting. Notably, many of these factors or marks have been reported to play an important role in iPSC reprogramming and pluripotency. BRD4 has been shown to play a crucial role in driving the late phase of iPSC reprogramming (Liu et al., 2014). Occupancy of the H3.3 variant has been attributed to maintaining MEF-specific identity during the early stages of pluripotency reprogramming. However, towards the fag end of reprogramming, they have been conducive to determine and maintaining pluripotent cell fate (Fang et al., 2018). Moreover, H3.3 is found to be critical for early development as its depletion results in early embryonic lethality (Jang et al., 2015). Ablation of H3.3 has been implicated to reduce the overall chromatin accessibility in ESC (Tafessu et al., 2023). Overall, H3.3 enrichment is crucial in pluripotent cell maintenance and early development. Also, enrichment of active chromatin marks, H3K27ac in enhancer and H3K36me3 in gene bodies of mouse embryonic stem cells are important for their maintenance (Kim et al., 2018; Mikkelsen et al., 2007). Altogether, we conclude that these chromatin accessibility factors drive reprogramming through mediating allelic bursting coordination of genes involved in iPSC reprogramming to fine-tune the appropriate dosage of these genes (Fig. 6). Broadly, our study provides fundamental insights into the implications of chromatin states in fine-tuning of allelic dosage of genes to orchestrate cell fate specification and extend strong support towards the role of chromatin state in mediating transcriptional bursting.

**Figure 6:**
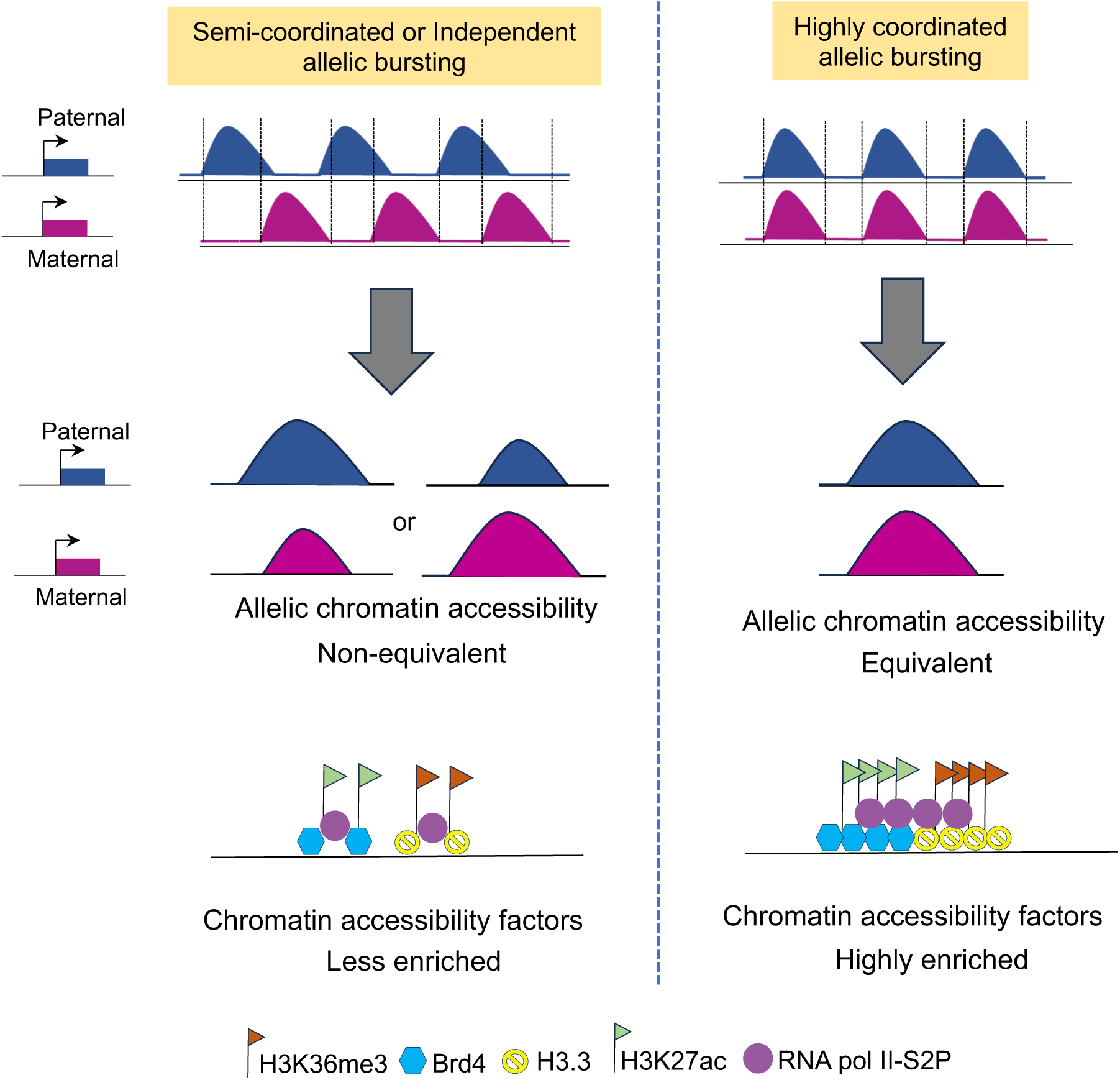
Chromatin state mediates allelic bursting coordination: Allelic chromatin accessibility is linked to allelic transcriptional bursting coordination. Genes with highly coordinated allelic bursting possess equivalent allelic chromatin accessibility. On the contrary, semi-coordinated or independent genes allelic chromatin accessibility differs. Importantly, highly coordinated genes are highly enriched with chromatin accessibility regulators.

## Materials and Method

### Data acquisition

Single-cell RNA-seq, ATAC-seq dataset for MEF to iPSC reprogramming were retrieved from Gene Expression Omnibus under the following accessions: GSE153846 and GSE153844 respectively (Talon et al., 2021). The ChIP-seq datasets used for MEF and iPSC were retrieved from GSE87037 (Aldiri et al., 2017), GSE99592 (Fang et al., 2018), GSE33823 (Yildirim et al., 2012) and GSE90893 (Chronis et al., 2017).

### Allele-specific expression and burst kinetics analysis

To obtain allelic read counts from scRNA-seq data, we performed allele-specific expression analysis following the pipeline as reported previously (Mandal et al., 2020; Naik et al., 2021, 2022). Briefly, we first constructed in silico CAST/EiJ and 129S1 specific parental genome by incorporating CAST/EiJ or 129S1 specific SNPs into the mm10 genome using variant calling file (VCF) tool (Danecek et al., 2011). VCF was downloaded from the mouse genome project (https://www.sanger.ac.uk/science/data/mouse-genomes-project). Next, we aligned RNA-seq reads into both parental genomes using STAR aligner (STAR-2.7.10a), allowing no multi-mapped reads (--outFilterMultimapNmax 1). We filtered out those genes for allele specific read counts which had at least 2 informative SNPs and minimum 3 reads per SNP site. We took an average across SNPs to get gene level allelic read counts. We normalized the allelic read counts by RPKM (reads per kilobase million). Since, low expressed genes are dropout prone, we removed low expressed genes from our analysis to avoid potential dropout effect (Kim et al., 2015; Santoni et al., 2017; Zhao et al., 2017). We considered only those genes, which shown minimum 25 cell expression and mean 10 RPKM across the cells. Allelic ratio was calculated individually for each gene using the formula = (129S1/CAST reads) ÷ (129S1+CAST reads). A gene was considered monoallelic if at least 95% of the allelic reads came from only one allele. We performed genome-wide allele-specific burst kinetics analysis using SCALE (Jiang et al., 2017). In brief, SCALE relies on Empirical Bayes Framework, which first classifies the genes into: monoallelic, biallelic and silent based on the allele-specific read counts and deduces the allelic burst kinetics based on the two-state model of transcription. It infers different burst kinetics parameters at the allelic level, such as burst frequency (Kon) and burst size (S/Koff), as described in the results. We excluded X-linked genes for our burst kinetics analysis.

### Gene ontology

Functional enrichment of different classes of genes was profiled using g:GOSt from gProfiler (https://biit.cs.ut.ee/gprofiler_archive3/e102_eg49_p15/gost) with Benjamini–Hochberg FDR and selected the biological process having FDR < 0.05 from GO: BP (Raudvere et al., 2019).

### Allelic ATAC-seq analysis

For allele-specific ATAC-seq analysis, first we created an ‘N-masked reference genome mm10’ through substituting strain-specific (129S1/SvImJ and CAST/EiJ) SNP position with ‘N’ using SNPsplit_genome_preparation (0.5.0) (F and SR, 2016) . Strain specific SNPs were obtained as described above. Next, reads for all samples were mapped to this N-masked genome using Bowtie2 (B and SL, 2012). We removed duplicate reads and mitochondrial reads from our analysis. SNPsplit was then used to create allele-specific BAM files by segregating the aligned reads into two distinct alleles (129S1/SvImJ and CAST/EiJ). Bigwig files were generated from these allelic BAM files using deepTools (version 3.5.1) function bamCoverage (--binSize 100 --smoothLength 500 --normalizeUsing RPGC) (Ramírez et al., 2016). Next, we performed enrichment analysis over the TSS region using deepTools.

### ChIP-seq analysis

For analysis of chromatin mark enrichment in MEF and iPSC in different categories of genes, we analysed available ChIP-seq data. For ChIP-seq analysis, the reads were mapped to mm10 genome using Bowtie2 (B and SL, 2012). We removed duplicate reads, mitochondrial reads from our analysis. Input normalized bigwig files were generated from the mapped BAM files and enrichment over the TSS and gene body region was plotted using deepTools (version 3.5.1).

### Quantification and statistical analysis

All statistical analysis was performed using the R software (https://www.R-project.org/). Mann–Whitney two-sided U test was used for statistical significance analysis and p values < 0.05 was considered as significant. For correlation analysis, Pearson test was used.

### Author’s Contribution

SG conceptualized, supervised, and acquired the funding for the study. Parichitran, HCN, performed experiments and analysis. AJN, HCN, Parichitran and LS helped in writing, conceptualization and interpretation. The final manuscript was edited and approved by all the authors.

## Acknowledgments

This study is supported by DBT grant (BT/PR30399/BRB/10/1746/2018), DST-SERB (CRG/2019/003067), DBT-Ramalingaswamy fellowship (BT/RLF/Re-entry/05/2016) and Infosys Young Investigator grant award to SG. AJN acknowledge Indian Institute of Science (IISc), Bangalore for the fellowship. LS acknowledge DST Women scientist award.

