## Supplementary Figure for "Chromatin states contribute to coordinated allelic transcriptional bursting to drive iPSC reprogramming"

### Supplementary figures

A

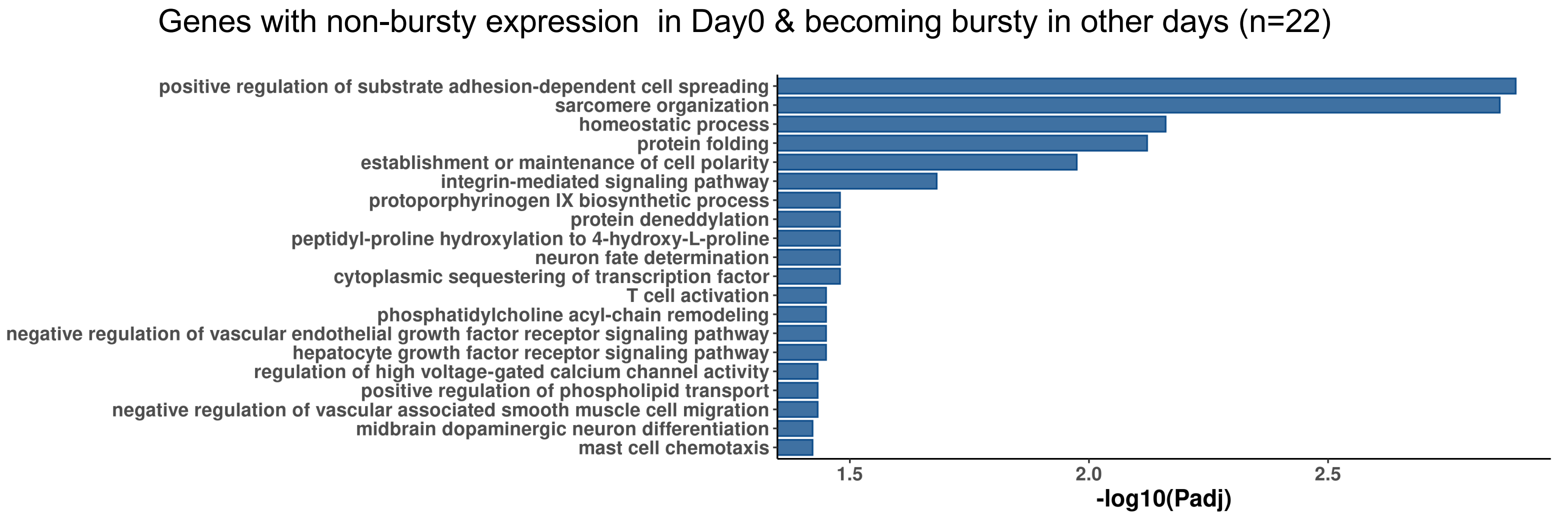


B

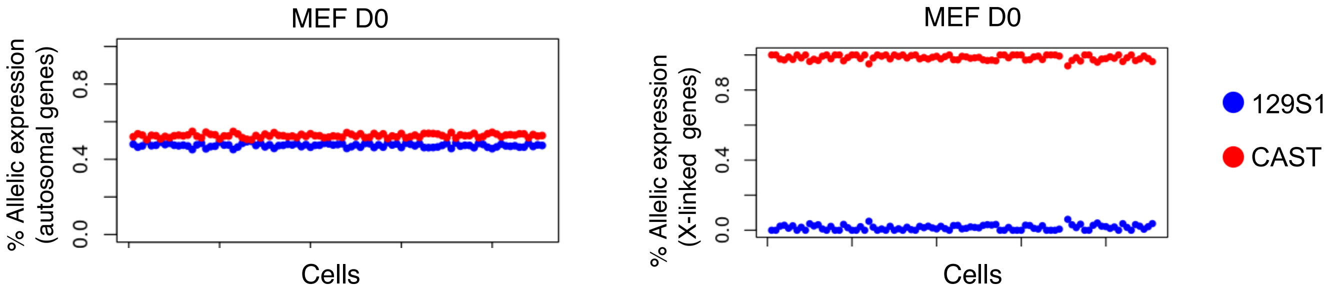


**Fig. S1, related to Fig. 1:** (A) Gene ontology analysis. (B) Plots representing allele-specific expression of autosomal and X-linked genes.

Fig. S2

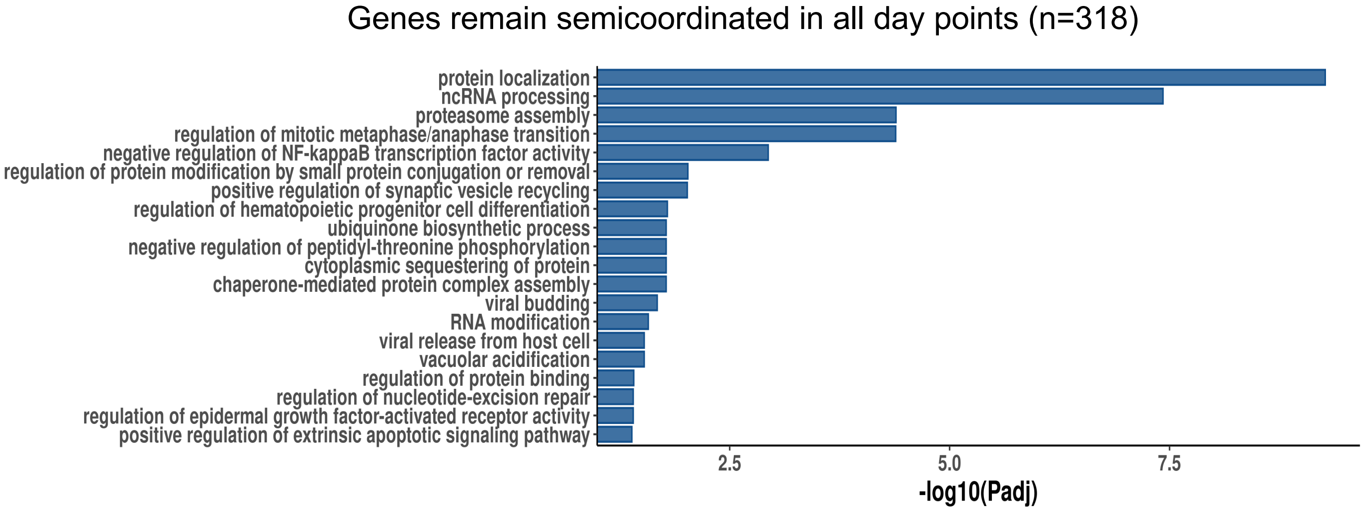


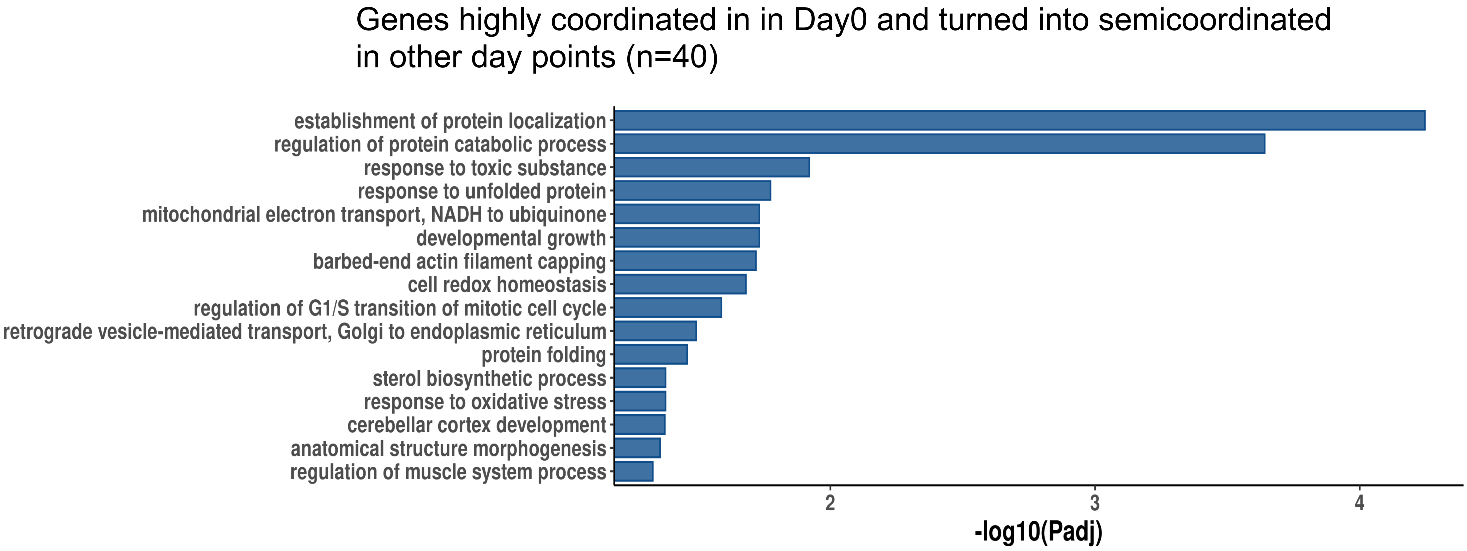


**Figure S2, related to Fig. 3:** Gene ontology (GO) enrichment analysis of genes (n=318) that remained semi-coordinated in all day points (top) and highly coordinated genes (n=40) in day 0 MEF that became semi-coordinated in other stages.

Fig. S3

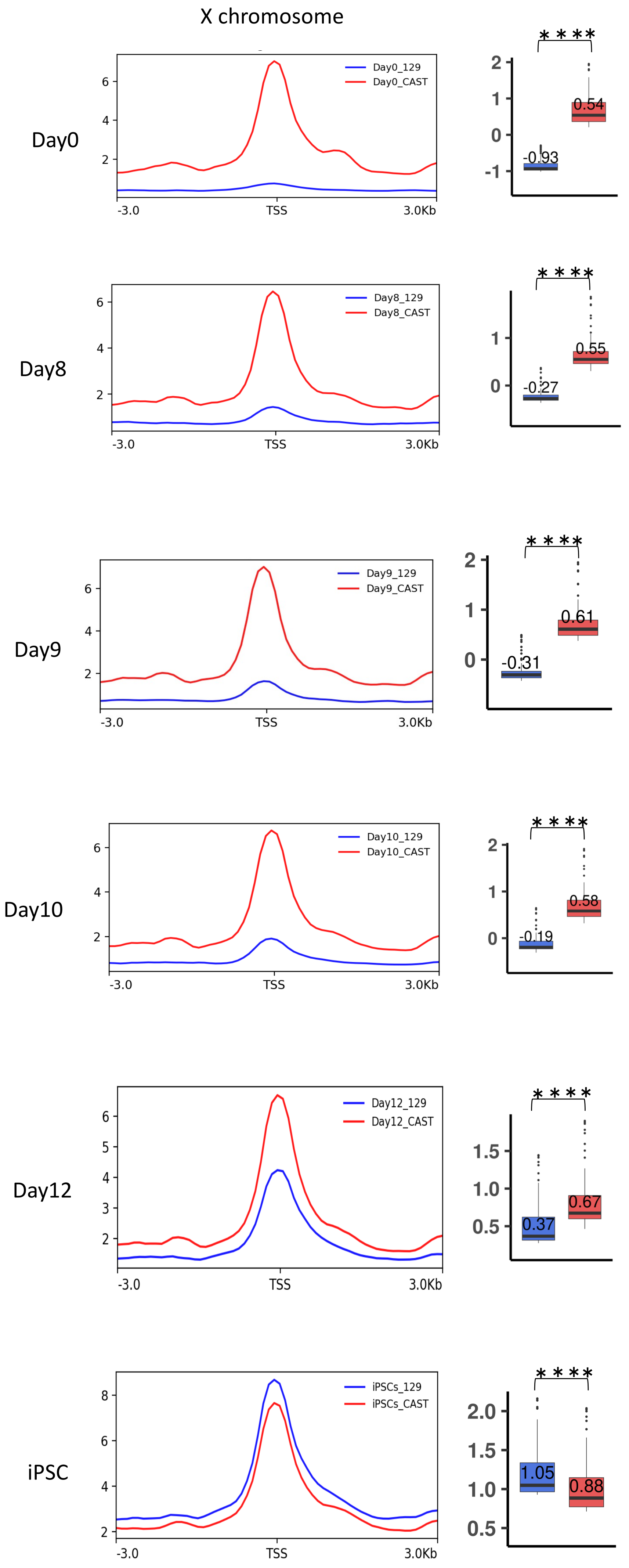
X chromosome
Day0
Day0_129
Day0_CAST
-3.0
TSS
3.0Kb
****
-0.93
0.54
Day8
Day8_129
Day8_CAST
-0.27
0.55
Day9
Day9_129
Day9_CAST
-0.31
0.61
Day10
Day10_129
Day10_CAST
-0.19
0.58
Day12
Day12_129
Day12_CAST
0.37
0.67
iPSC
iPSCs_129
iPSCs_CAST
1.05
0.88

**Figure S3, related to Fig. 4: Allelic chromatin accessibility for X-linked genes.** Quantification of enrichment of allelic accessibility across 3 kb upstream and 3 kb downstream of TSS of X-linked genes in all day-points of reprogramming. In the boxplots, the line inside of each box signifies median value whereas the edges of each box denote 25% and 75% of the datasets (Wilcoxon Rank Test: p *value* < *0.0001*; ****).

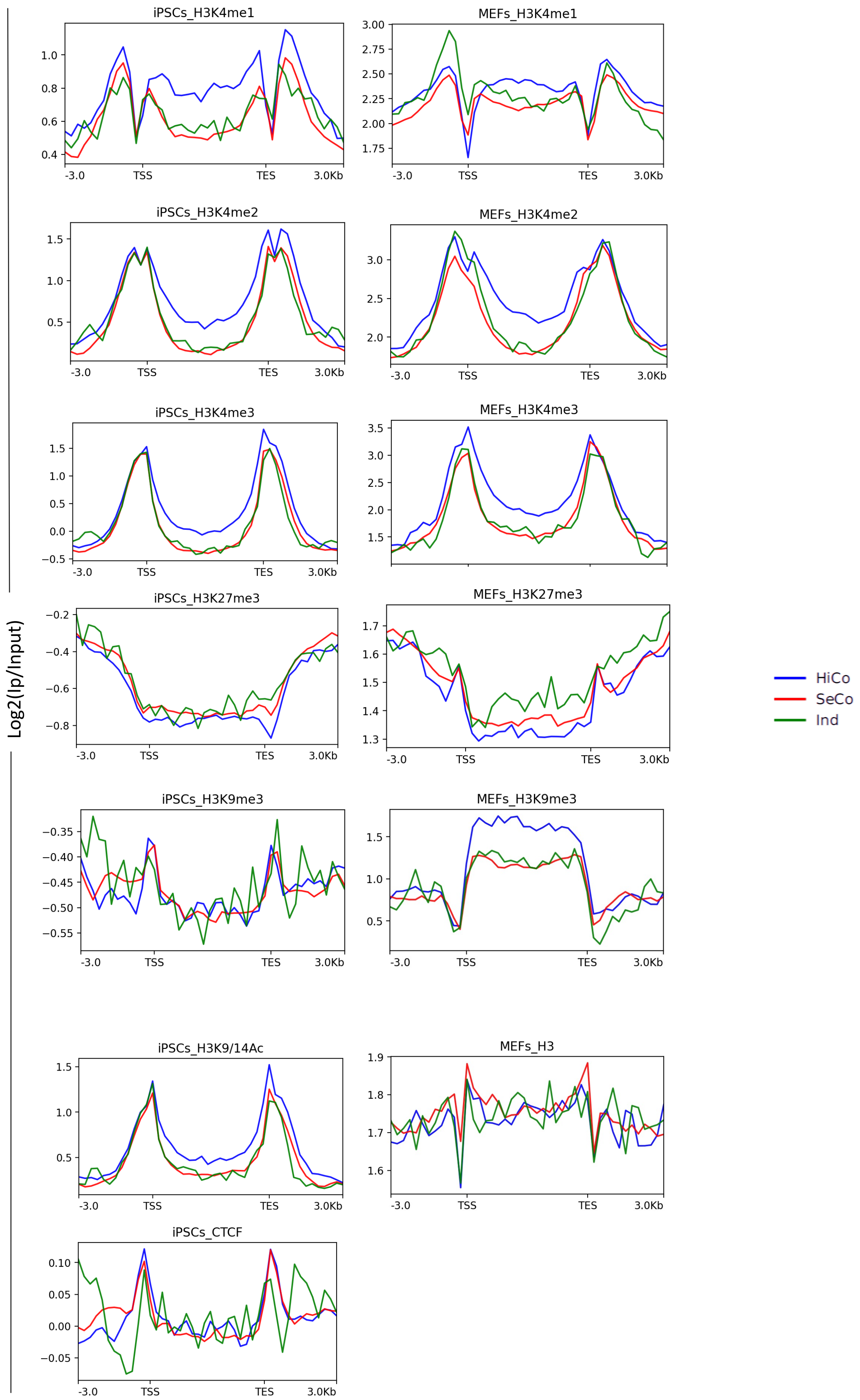


**Figure S4, related to Fig. 5:** Comparison of enrichment of different chromatin accessibility related factors in the gene body and TSS of highly coordinated, semi-coordinated and independent genes in MEF and iPSC.
